# Biogenic plate-like guanine crystals form via templated nucleation of thin crystal leaflets on amyloid scaffolds

**DOI:** 10.1101/2022.09.29.510168

**Authors:** Zohar Eyal, Rachael Deis, Neta Varsano, Nili Dezorella, Katya Rechav, Lothar Houben, Dvir Gur

## Abstract

Controlling the morphology of crystalline materials is challenging, as crystals have a strong tendency towards thermodynamically stable structures. Yet, organisms form crystals with distinct morphologies, such as the plate-like guanine crystals produced by many terrestrial and aquatic species for light manipulation. Regulation of crystal morphogenesis was hypothesized to entail physical growth restriction by the surrounding membrane, combined with fine-tuned interactions between organic molecules and the growing crystal. Using cryo electron tomography of developing zebrafish larvae, we found that guanine crystals form via templated nucleation of thin leaflets on preassembled scaffolds made of 20-nm-thick amyloid fibers. These leaflets then merge and coalesce into a single plate-like crystal. Our findings provide new insights into how organisms control the morphology and, thereby, the optical properties of crystals.

---

The nanoscale morphologies of crystalline materials determine their optical, electrical and mechanical properties and, thus, their potential application(*1–4*). In nature, many organisms use molecular crystals for their function, which they form out of small organic molecules under ambient conditions(*5–8*). A prominent example are the thin plate-like guanine crystals, utilized by a huge variety of terrestrial and aquatic organisms for diverse functions such as vision, camouflage, body temperature regulation, and kin recognition(*5, 6, 9*). Guanine crystals are constructed from H-bonded molecular layers that are stacked one of top of the other by π-stacking(*10*). Plate-like guanine crystals expose the extremely high, in-plane refractive index (n=1.83) of light, thereby allowing the construction of highly effective and versatile photonic arrays (**Fig. 1 A and B**)(*11*). However, the inherent thermodynamic properties of the crystalline lattice creates a strong tendency toward forming specific low-energy prismatic structures(*6, 12, 13*). Overriding this intrinsic tendency of the crystals to grow as prisms requires extensive biological intervention to selectively impede the growth along the π-stacking direction (**Fig. 1 A and B**). Yet, the mechanism that exerts such tight and extensive control over crystal morphology has remained a mystery.

**Figure 1.**
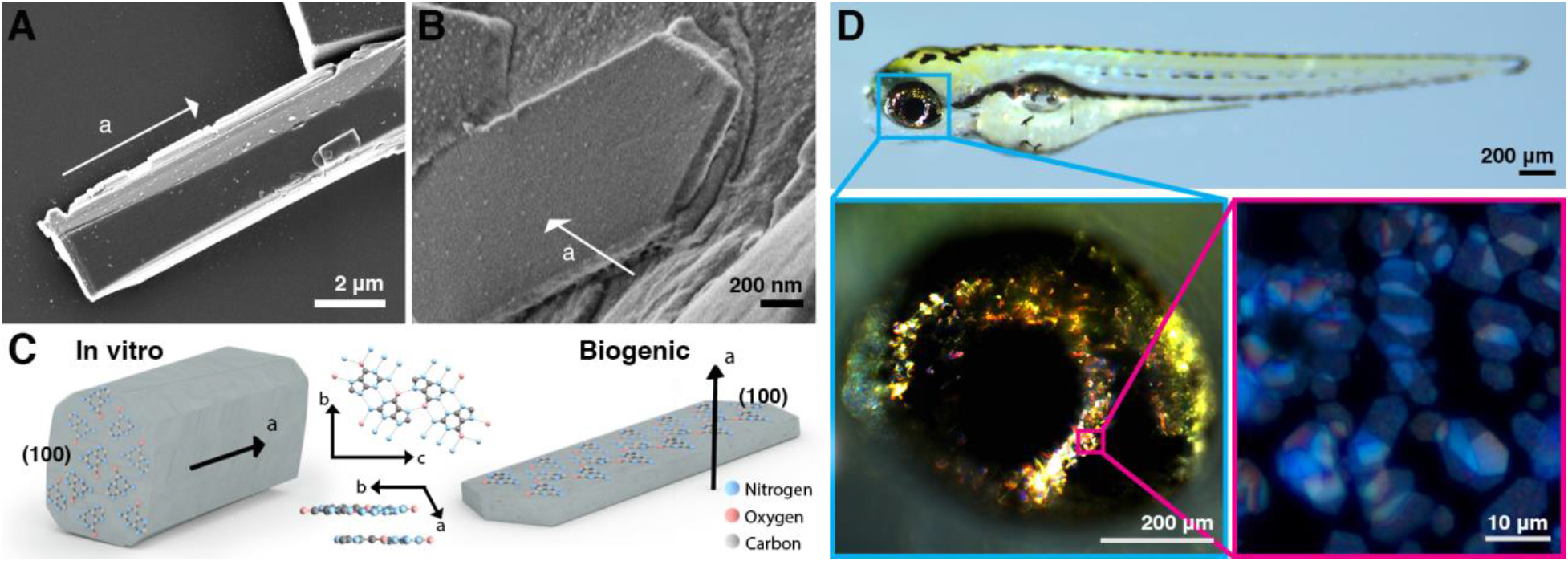
Synthetic and biogenic guanine crystals have distinct morphologies. (**A**) A synthetic guanine crystal with a prismatic morphology, where the (100) crystallographic plane is the fastest growing direction. (**B**) A biogenic plate-like biogenic guanine crystal isolated from a zebrafish eye. (**C**) Schematic illustration of both synthetic and biogenic crystals in which the (100) crystallographic plane, constructed from H-bonded molecular layers, is shown. (**D**) A zebrafish larva at 5 days post-fertilization (dpf) that contains guanine crystals in its eyes and skin. Insets show higher magnifications of the entire eye (cyan) and of the iridophores and the crystals within them (pink). A and B are SEM micrographs and D shows incident light images.

Specialized guanine crystal-forming cells, also known as iridophores, produce crystals within membrane-bound organelles termed iridosomes(*14, 15*), where exquisite control over crystal size, shape and assembly far exceeds the synthetic state-of-the-art in materials science and solid-state chemistry(*6, 16*). Iridophores are often compared to melanophores, cells that absorb light rather than reflect it(*17*). Studies on melanophores indicate that their melanin-producing organelles, called melanosomes, are derived from endosomal compartments, and belong to the lysosome-related organelles (LROs) family(*18*). However, much less is known about non-melanosomal pigment organelles and the regulation of the processes occurring within them(*19, 20*). Over the years, several mechanisms that control biocrystal morphogenesis have been suggested, including the involvement of different biochemicals that interact with the growing crystal or that get incorporated into its lattice(*8, 21, 22*). These biomolecules were postulated to interact with the growing crystals stereochemically and thereby create local kinetic conditions that allow the formation of diverse crystal morphologies. However, studies of the biochemical composition of guanine crystals with different morphologies from a variety of organisms, suggest that the small molecule composition of crystals did not correlate with their shape (*23*). In spiders, prismatic crystals were proposed to form on sheets within a crystal vesicle(*24*) and to be comprised of 25 nm-thick crystal lamellas stacks (*25*). Studies on inorganic biocrystals suggest that crystal morphogenesis is controlled by physical confinement of growth by the delimiting membrane of the organelle(*26*). The rationale here being that confinement allows for growth rate manipulations, which are sufficient to produce a variety of complex shapes.

To elucidate the morphogenesis of guanine plate-like crystals and the underlying mechanism, we investigated iridophores in the zebrafish (*Danio rerio*) model organism, which uses guanine crystals in their eyes as both a light barrier, and for enhancing vision sensitivity under low light conditions (**Fig. 1C**)(*27, 28*). Synchronized crystal growth in the zebrafish larva iridophores starts as early as 44 hours post-fertilization (hpf), making it an ideal system to follow the early stages of crystal formation(*29*). To study the early events of crystal formation in zebrafish larvae eyes in their native hydrated state, we used cryogenic scanning electron microscopy (cryo-SEM) (**Fig. 2 A-D**). By using this technique we imaged iridophores adjacent to melanophores that, in turn, are adjacent to photoreceptors in the retina (**Fig. 2 A-C**). In adult fish, mature iridophores are packed with dozens of membrane-bound plate-like crystals (**Fig. S1**). At earlier developmental stages, the plate-like crystals were composed of very thin leaflets, a few nanometers in thickness, representing only 5-20 planner molecular stacks of guanine sheets (**Fig. 2D**). During subsequent maturation, the leaflets expanded and eventually coalesced into a single crystal (**Fig. 2D**).

**Figure 2.**
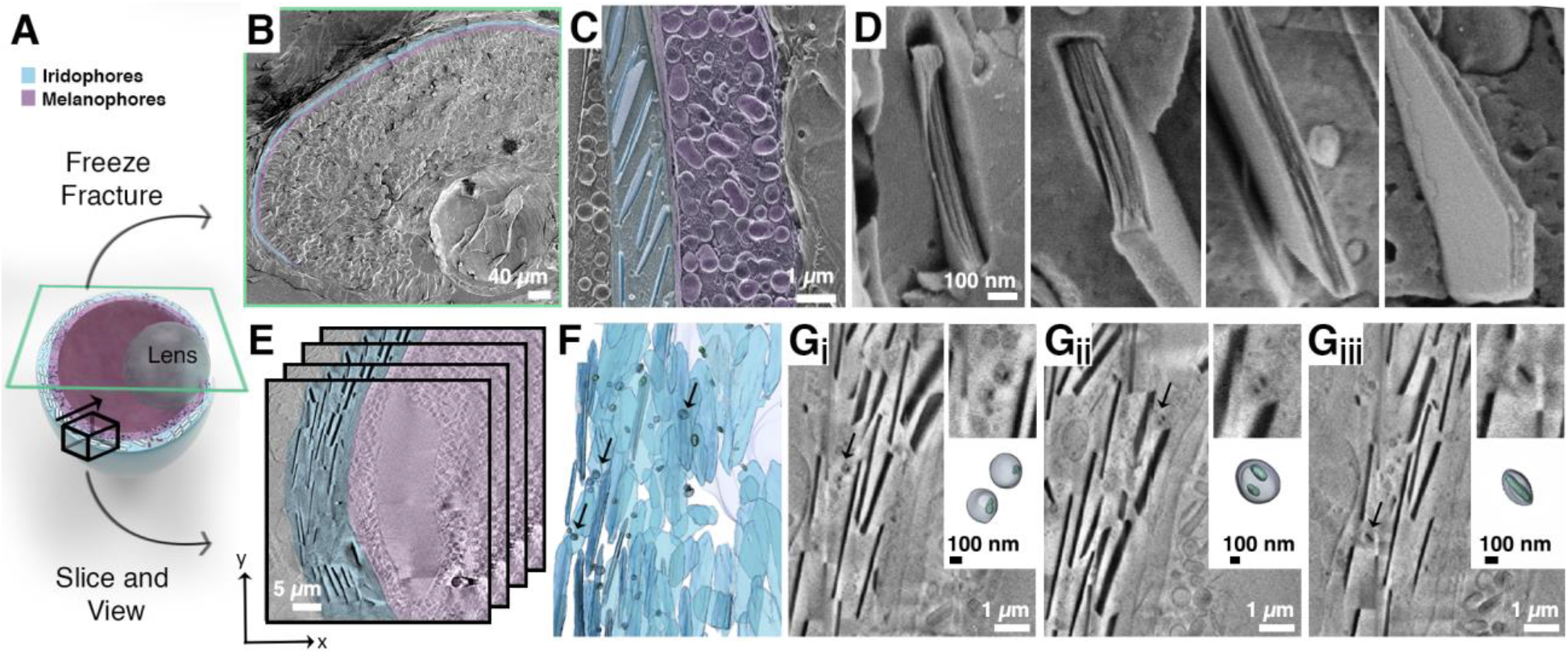
Early iridosomes contain several crystal leaflets that later coalesce into a single plate-like crystal. (**A**) An illustration of the zebrafish larva eye showing the iridophore (blue) and melanophore layers (purple), next to the lens. The anatomical location of the sections and areas that were taken for cryo-SEM (B-D) and cryo-FIB-SEM (E-G) are marked by a green rectangle and a black box, respectively. (**B-D**) Cryo-SEM images of freeze-fractured surfaces of a zebrafish larva eye and the iridophores within it. (B) A low magnification of the eye surface, where the iridophore and melanophore layers are pseudo-colored in blue and purple, respectively. (C) A close-up view of an iridophore next to a melanophore both pseudo-colored as in (C). (D) Iridosomes at different maturation stages. The plate-like crystals are initially composed of very thin leaflets, which grow and eventually coalesce into a single crystal. (**E-G**) Cryo-FIB-SEM images and 3D representations of segmented milled sections from a freeze-fractured zebrafish larva eye. (E) An illustration showing the stack of images of consecutive volumetric sections, in which iridophore and melanophore layers are pseudo-colored in blue and purple, respectively. (F) A 3D representation of an eye iridophore, where both mature elongated iridosomes and round developing iridosomes are marked in blue and the cell nucleus is marked in gray. The early 200-300 nm long iridosomes shown in (G_i_-_Giii_) are marked with black arrows. (Gi-G_iii_) Cryo-FIB-SEM micrographs of different areas within the iridophore shown in (F). Black arrows mark the iridosomes whose 3D reconstructions are shown in the insets.

Capturing the initial transient stages of crystal formation is challenging and requires obtaining volumetric 3D data, which allow examination of the entire volume of the iridosomes in their native state. Therefore, we used cryogenic focused ion beam SEM imaging (cryo-FIB-SEM) to study early iridophores in situ, inside the zebrafish larva eyes (**Fig. 2 E-G**). In cryo-FIB-SEM, guanine crystals appear as dark contrast objects, due to their surface potential. Using this approach, we found small, ~150-300 nm round iridosomes next to more mature, elongated iridosomes, which were already several micrometers in length (**Fig. 2 F and G**). Surprisingly, the small and round iridosomes contained thin, well-developed crystals (**Fig. 2 F and G and Fig. S2**). As crystal growth advanced, iridosomes gradually acquired the typical elliptical shape of the mature organelle, in which the membrane tightly engulfs the elongated crystals. In the smallest and most immature iridosomes, we observed distinct thin leaflets that did not appear to fill the entire volume of the iridosome. In certain cases, multiple thin leaflets were present in the same iridosome (**Fig. 2G**).

Since the early stages of crystal formation occur at the nanometric scale, they are barely visible using cryo-SEM and cryo-FIB-SEM modalities. Thus, we utilized high-resolution cryo electron tomography (cryoET) to investigate crystal formation in 3D at different developmental stages (**Fig. 3**). Notably, in this approach the intact membranes are clearly visible, which allows studying the interface between the membrane and developing crystal and the role of confinement in this process. To image iridosomes from different developmental stages, we plunge-freezed iridophores that were isolated from *pnp4a:palmmCherry* transgenic zebrafish larvae using fluorescence-activated cell sorting (FACS) (**Fig. S3**). In iridophores that were isolated from ~72 hpf larvae, we observed iridosomes with two main types of crystal organization. Crystals that were positioned with their (100) facet either face-on (**Fig. 3 A, D, G**) or edge-on (**Fig. 3 C, E**, **F, H**) with respect to the electron beam. In all observed iridosomes, the initial crystals formed within the lumen of the organelle, with no apparent contact with the surrounding bilayer membrane (**Fig. 3 A and C**). The elongated crystals then continued to grow towards the surrounding membrane until they eventually made contact, and started pushing against it (**Fig. 3D**). At this point, additional crystal growth resulted in deformation and elongation of the surrounding membrane (**Fig. 3G**), until both the crystal and the iridosome membrane assumed their final elongated morphology (**Fig. S3**).

**Figure 3.**
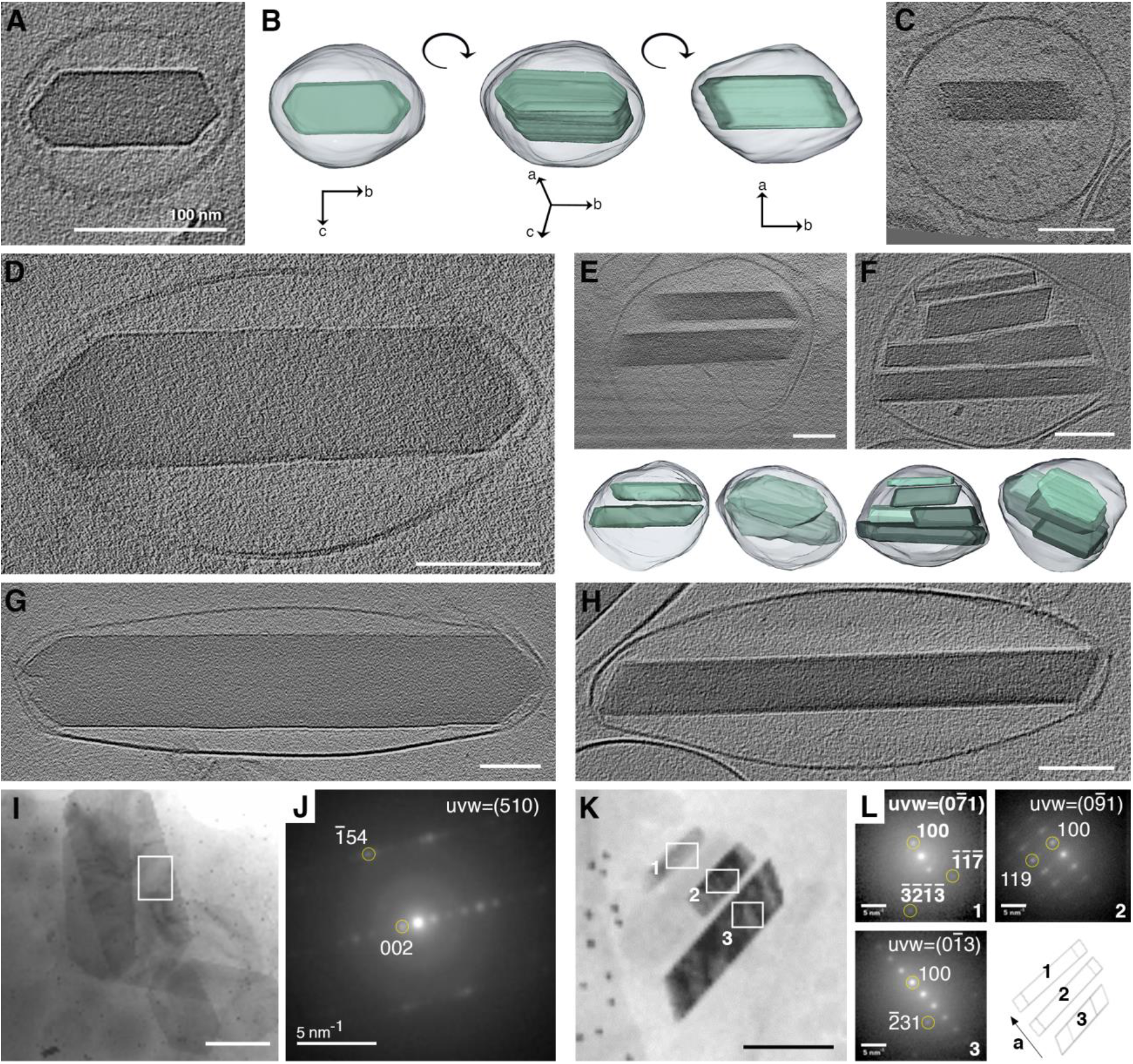
High-resolution cryo-TEM shows the morphological sequence of (100) crystal plates within iridosomes. (**A, C-F**) Cryo-ET reconstructions of iridophores, isolated from zebrafish larvae at different developmental stages, show crystals either face-on (A,D,G) or edge-on (C,E,F,H) with respect to the electron beam. (**B**) A 3D representation of a crystal-containing iridosome from different angles, emphasizing the different orientations. Bottom panels in (E) and (F) show 3D representations of leaflet-containing iridosomes. (**I,K**) Cryo-4D-STEM bright field images of iridosomes with crystals orientated face-on (I) or edge-on (K) with respect to the beam. (**J**) Electron diffractions taken from the area marked by a white rectangle in (I). (**L**) Electron diffractions taken from the area marked by a white rectangle in (K), and an illustration showing the orientation of the *a* axes of the crystals (bottom right corner).

In iridosomes where the plate-like crystals were orientated edge-on, we often observed 2-8 distinct crystals leaflets within the same organelle (**Fig. 3E-F, and supporting movie 1**). To correlate the morphological information with the crystallographic features of the crystals observed, we investigated iridosomes using cryogenic 4D scanning transmission electron microscopy (cryo-4D-STEM), exploiting its scanning nanobeam electron diffraction (NBED) abilities to collect an electron diffraction pattern from every point the beam raster traverses (**Fig. 3 I-L**). We found that crystal leaflets that were oriented face-on exhibited a diffraction pattern that correlated to the (002) crystallographic plane (**Fig. 3 I-J**). Whereas the diffraction pattern observed of leaflets that assumed the perpendicular orientation correlated to the (100) crystallographic plane (**Fig. 3 K-L**). These results strongly indicate that the very initial leaflets are already formed as (100) crystal plates. In developing iridosomes that contained multiple crystal leaflets, we often observed that the leaflets were not fully aligned (**Fig. 3 E-F, Fig. S4**). However, when we used NBED to map the orientation of crystals in more developed iridosomes, we found that the crystal leaflets were arranged such that their *a* axes were almost completely parallel (**Fig. 3L**). Remarkably, in mature iridosomes, the leaflets coalesced into a single crystal with no obvious remnants of the initial leaflets.

Our observations that the initial crystals form in the lumen of the iridosome, with no apparent contact with the surrounding membrane, already as thin (100) crystal plates excluded significant contribution of physical confinement to the obtained plate-like morphology. We therefore analyzed iridophores isolated from younger zebrafish larvae (44-48 hpf) to investigate the early nucleation events during crystal leaflet formation **(Fig. 4**). Intriguingly, we found that the iridosomes in these very early cells contained multiple parallel fibers, approximately 20 nm in diameter and 200-400 nm in length, running across them (**Fig. 4A and Fig. S5**). Imaging of more developed iridosomes revealed small crystals in close contact with the preassembled fiber scaffold (**Fig. 4B**). Remarkably, these initial crystals already formed as thin leaflets, assuming the typical (100) plate-like morphology. Staining with Thioflavin T (ThT), a well-established amyloid marker(*30*), indicated that these fiber scaffolds are proteins, aggregated as amyloid fibers (**Fig. S6**).

**Figure 4:**
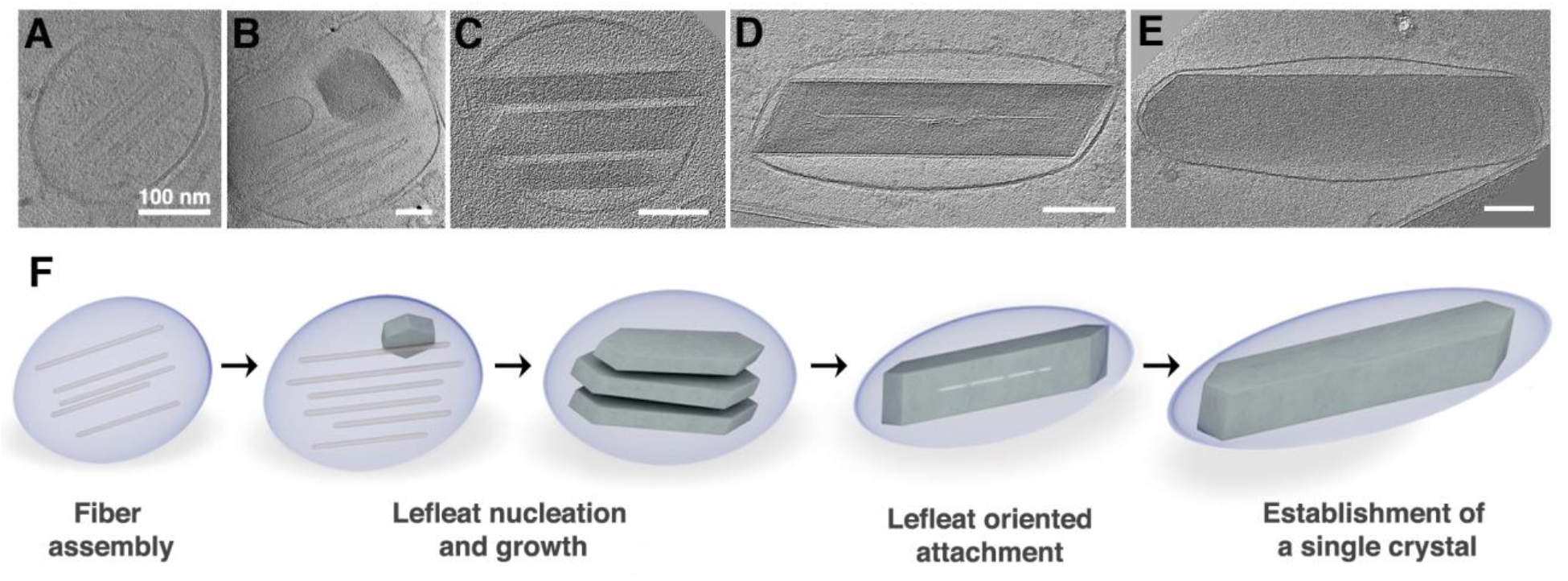
Biogenic plate-like guanine crystals form via templated nucleation of (100) crystal leaflets on preassembled protein fibers. **(A-E)** CryoET reconstructions of iridophores isolated from zebrafish larvae at different maturation stages. (A) An early iridosome showing a scaffold of parallel protein fibers. (B) A more developed iridosome, in which the initial nucleation of (100) crystal leaflet on a protein scaffold is taking place. (C) Several individual leaflets, imaged edge-on, with their (100) crystal plane parallel to each other. (D) The leaflets have almost completely coalesced into a single crystal. (E) A mature iridosome, where the leaflets have completely merged into a single crystal, with no obvious traces of the initial leaflets **(F)** A schematic illustration of the proposed crystallization mechanism.

The formation of thin plate-like guanine crystals requires overriding the strong intrinsic tendency of crystals to grow as prisms and to preferentially express the thermodynamically disfavored (100) face (parallel to the H-bonded plane)(*10*). Recent studies suggest that the control over morphology of certain biocrystals can be obtained via confinement, i.e., a physical block of growth by the delimiting membrane(*26*). However, we found that the initial crystal formation within the iridosome lumen occurs at a considerable distance from the delimiting membrane. Only later in development, the elongating crystals reach the membrane, eventually making contact and pushing against it. Based on our high-resolution cryoET of iridophores at different developmental stages, we propose that the formation of plate-like crystals occurs via templated nucleation of thin leaflets on preassembled amyloid fiber scaffolds. This process begins with the assembly of the protein scaffolds inside a ~100 nm iridosome (**Fig. 4A**). Then, nucleation of distinct (100) crystal plates takes place on the fibers (**Fig. 4B**). As development proceeds, the number and size of crystal leaflets increases (**Fig. 4C**), followed by their alignment and gradual oriented attachment (**Fig. 4D**), eventually culminating in the formation of a single plate-like guanine crystal (**Fig. 4E**). Figure 4F schematically illustrates this sequence of events. The preassembled amyloid fiber scaffolds likely drive the crystal growth along the H-bonded direction through molecular recognition. This recognition could occur either at the level of the basic building blocks, i.e., interactions between specific amino acids and the guanine molecules, or at a macromolecular level, promoting the interaction of hydrophobic domains in the fiber with the planar sheets of H-bonded guanine layers. In organelles of other pigment cells, namely melanosomes, it was shown that fibrillar sheets of PMEL, a non-pathogenic amyloid protein, serve as a template for melanin polymerization and synthesis, and that lack of PMEL results in different degrees of hypopigmentation(*31, 32*). Identification of the protein which forms the amyloid scaffolds in iridosomes, and uncovering its structure could further elucidate the mechanism of controlled crystal formation, shedding new light on the interactions between proteins and molecular crystals and on the mechanisms that control crystal nucleation and growth.

## Acknowledgments

This work was supported by the Israel Science Foundation grant 691/22. We thank Dr. Smadar Zeidman and Dr. Eyal Shimoni for assistance with the conventional TEM preparations and data collection. Additionally, we thank Prof. Lia Addadi for helpful scientific discussions. We thank Dr. Tali Lerer-Goldshtein and Dr. Anna Gorelick-Ashkenazi for their great support with developing the project experimental approach. Electron microscopy studies were conducted at the Irving and Cherna Moskowitz Center for Nano and Bio-Nano Imaging at the Weizmann Institute of Science.

## Notes

### Competing Interest Statement

The authors have declared no competing interest.

